# DNA sequence quantitatively encodes CTCF-binding affinity at genome scale

**DOI:** 10.64898/2026.01.05.696797

**Authors:** Zihang Yin, Yefan Wang, Guojun Hou, Yuqiao Chen, Huaijin Feng, Jin Zhu, Yufan Xie, Ruoxi Wang, Zhiyu Zhang, Tao Wu, Haiyan Huang, Sheila Q. Xie, Wuming Wang, Wenchao Gu, Qiang Wu, Nan Shen, Ya Guo

## Abstract

CTCF is a central architectural protein that shapes 3D genome organization through sequence-specific DNA binding, but how DNA sequence quantitatively determines CTCF-binding strength remains poorly understood. Progress has been hampered by the lack of large, high-quality measurements of binding affinity. Here, we experimentally determine *in vitro* CTCF-binding affinity for 276,765 DNA sequences derived from the human genome, generating a comprehensive quantitative landscape of CTCF-DNA interactions. Leveraging these data, we develop DeepCTCF, a deep learning model that predicts CTCF-binding strength directly from DNA sequence and enables quantitative interpretation of CTCF motif grammar. Using this framework, we systematically dissect how specific sequence features modulate CTCF-binding affinity and generate quantitative predictions for disease-associated variants that alter CTCF binding. Together, this study defines general principles by which DNA sequence encodes CTCF-binding affinity and provides a quantitative framework for interpreting regulatory sequence variation.

**Teaser:** Mapping how DNA encodes CTCF binding reveals quantitative rules across over 1‰ of the human genome.

## Introduction

Interphase chromosomes are folded into topologically associating domains (TADs) (*1, 2*), predominantly mediated by CCCTC-binding factor (CTCF) through cohesin-driven loop extrusion (*3-5*). CTCF/cohesin-mediated loops are dominantly formed between CTCF-binding sites in a convergent orientation (*6-9*). Thus, 3D-genome architecture can be encoded by linear genomic sequences (*9*). The CTCF-based interaction networks are mainly defined by three variables: location, orientation, and affinity of CTCF binding. Compared to the location and orientation, CTCF-binding affinity exhibits substantial variability and remains largely underexplored.

CTCF is essential for early embryo development (*10*), as well as for the normal development of cardiomyocytes and limbs (*11, 12*). CTCF also plays critical roles in transcriptional insulation (*13*), maintenance of genomic imprinting (*14-16*), control of V(D)J recombination (*17*), promoter choice of clustered protocadherin genes (*6, 18, 19*), inhibition of antisense transcription (*20*), and regulation of alternative splicing (*21*). These functions of CTCF in the control of gene expression are contextualized by its interplay with DNA. Using its tandem 11-zinc finger (ZF) domains, CTCF can bind to DNA at the core motif-1 recognized by the ZFs 3-7, while additional ZFs are believed to stabilize the binding (*22-26*). Initial analysis identified 13,804 CTCF sites in the human genome (*27*). Later, 35,161 CTCF-binding positions were detected (*28*). Of these, 96.8% of sites had the core motif-1, while 8,899 sites contained a 9-mer upstream motif (motif-2), which was also independently identified (*26, 29*). 8,857 CTCF-binding events carried a downstream motif (motif-3), and some rare events (550) contained a different upstream motif (motif-2’) (*28*), consistently identified (*22, 26*). Despite these advancements, *in vivo* CTCF binding may be regulated by DNA methylation (*30*), chromatin remodeling complex imitation switch (ISWI) (*31-33*), and protein-RNA interactions (*34, 35*), such that *in vivo* occupancy reflects not only DNA sequence but also chromatin context and cofactors, leaving the intrinsic binding affinity encoded by DNA sequence unclear.

CTCF-binding disruption is increasingly recognized as a mechanism by which missense variants cause diseases (*36-38*). Moreover, 3D-chromatin organization can be manipulated by engineering CTCF-binding sites (*9, 39*). Quantitative prediction of CTCF binding can thus help identify CTCF-binding-altering variants, and enables the de novo design of CTCF-binding sites. Early position weight matrix (PWM) models estimate the changes in CTCF binding based on position frequency matrices, but often suffer from high false positive predictions due to neglected positional dependencies (*40, 41*). Deep-learning models have been developed to address these limitations. For example, DeepBind uses convolutional neural networks (CNN) to learn sequence-specific binding patterns directly from protein-DNA binding data (*42*). BPNet further improves predictive resolution by combining CNN networks with base-resolution datasets (*43*). However, the performance of existing methods for CTCF-binding prediction remains constrained, largely owing to the limited availability of large, high-quality datasets suitable for quantitative modeling.

Here, we have created a large, high-quality dataset that enables the training of an artificial intelligence (AI) based framework to directly predict CTCF-binding affinity. By integrating computational modeling with targeted experimental validation, we identify sequence-context features that quantitatively shape CTCF-binding affinity, representing a critical step toward the de novo design of CTCF-binding sites. Moreover, DeepCTCF provides quantitative estimation of how sequence variations alter CTCF binding, enabling the exploration of regulatory mechanisms and therapeutic insights.

## Results

### A quantitative landscape of CTCF-binding affinity

CTCF-binding sites defined by the chromatin immunoprecipitation (ChIP)-based assays typically span 19 to 42 bp (*27-29*). Given that the sequence space of a 42-nucleotide region exceeds 10^25^ possible combinations, it is technically infeasible to directly measure the binding strength of CTCF for all potential sequences. To overcome this limitation, we developed a high-throughput approach, MpEMSA-seq, that enables parallel quantification of CTCF-binding affinity for 276,765 unmodified DNA sequences *in vitro* (Fig. 1A).

**Fig. 1.**
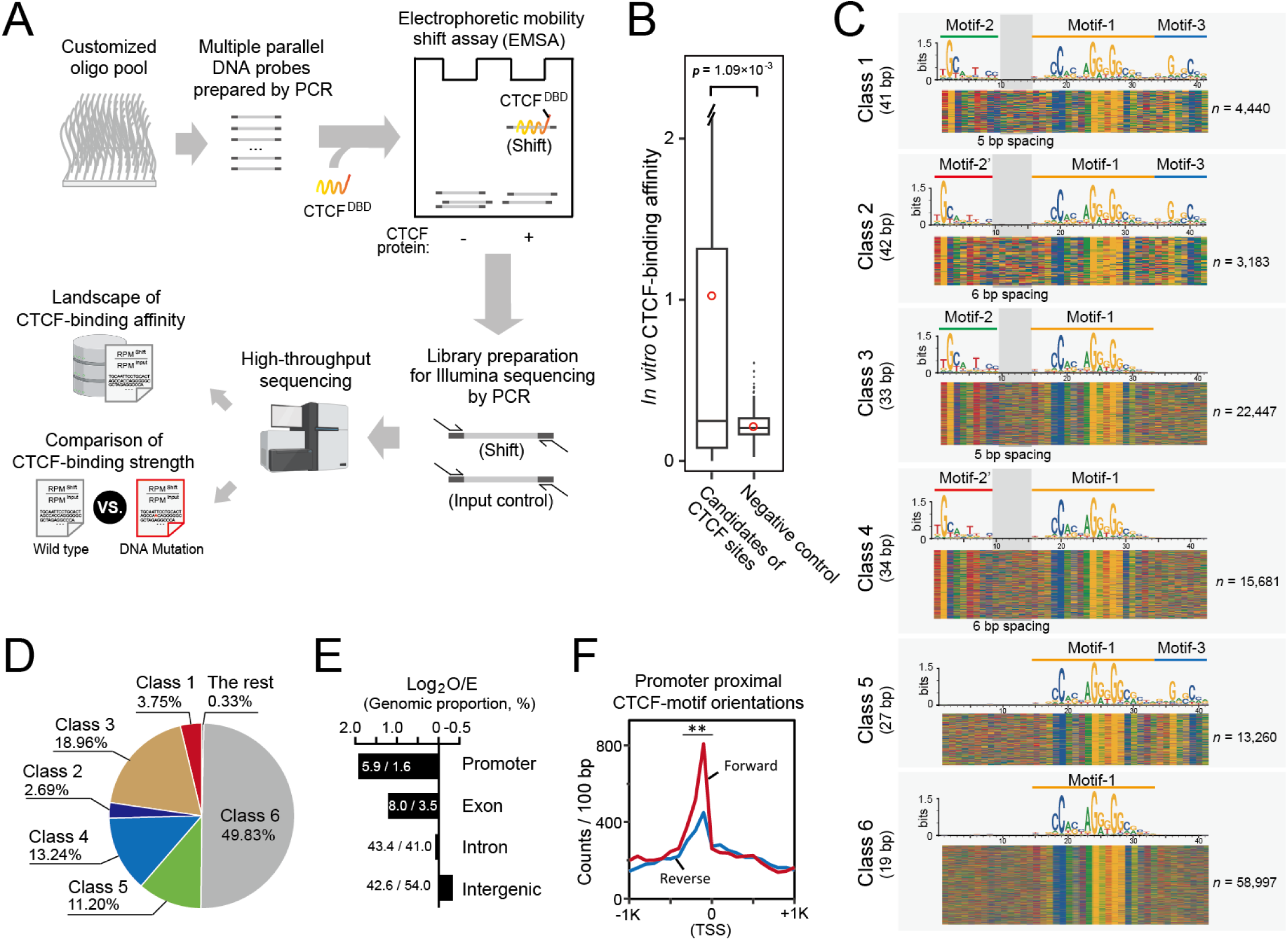
A quantitative landscape of CTCF-DNA binding affinity across the human genome. (**A**) Schematic of the MpEMSA-seq workflow, which combines multiple parallel electrophoretic mobility shift assays with high-throughput sequencing to quantify binding affinity between the CTCF DNA-binding domains (CTCF^DBD^) and 276,765 genomic DNA sequences. (**B**) Box plots comparing CTCF-binding affinity values for candidate CTCF sites and randomly generated sequences used as negative controls. Red circles indicate mean values. Boxes span the first to third quartiles, with horizontal lines indicating medians; whiskers extend to 1.5 ×the interquartile range (IQR), and points beyond the whiskers denote outliers. Statistical significance was assessed using the Mann-Whitney U test. (**C**) Sequence-based classification of identified CTCF-binding sites. Nearly all sites (99.7%) are assigned to six classes (classes 1-6) based on combinations of CTCF motifs (-1, -2, -2′, and -3). Each row represents a bound sequence, with color-coded nucleotides (A, green; T, red; G, yellow; C, blue) shown for 41-42 bp. (**D**) Relative proportions of the six classes of CTCF recognition sequences. (**E**) Enrichment of identified CTCF-binding sites across genomic annotations, shown as the log_2_ ratio of observed to expected counts in promoter, exon, intron, and intergenic regions. (**F**) Orientation bias of CTCF-binding sites relative to transcription start sites (TSS), quantified as counts per 100 bp for forward (red) and reverse (blue) orientations. ** *p* < 10^-15^, two-sample Poisson rate test.

We first evaluated the potential impact of adapter sequences on quantitative affinity measurements by testing three distinct adapter pairs (fig. S1A). The resulting affinity values showed strong concordance, with Spearman’s correlation coefficients ranging from 0.89 to 0.94. We then synthesized an oligonucleotide library comprising 276,765 candidate human CTCF-binding sequences flanked by adapter-1. These candidates were compiled from public ChIP datasets and *in silico* predictions (fig. S1B). Relative affinity values were calculated for each sequence. To identify CTCF recognition sequences that could be distinguished from nonspecific binding to random DNA, we applied two complementary statistical criteria. First, 124,223 sequences exhibited significantly higher affinity than 393 randomly generated control sequences based on a Mann-Whitney U test (*p* < 0.05). Second, 118,395 sequences showed average affinity values exceeding those of 95% of random sequences (Fig. 1B and table S1). Applying both criteria yielded a high-confidence set of 117,974 CTCF recognition sequences, collectively covering more than 1‰ of the human genome.

We next benchmarked MpEMSA-seq against a previously developed *in vitro* method (*44*). MpEMSA-seq identified approximately fourfold more CTCF recognition sequences than the recently reported GHT-SELEX approach, which captures genomic sequences directly bound by CTCF *in vitro* (fig. S1C). Notably, 91.1% of GHT-SELEX-identified CTCF sequences (29,920 of 32,853) were also detected by MpEMSA-seq. In addition, MpEMSA-seq identified 101,799 genomic loci not detected by GHT-SELEX. These uniquely identified sequences generally exhibited lower binding affinity (fig. S1D), indicating that MpEMSA-seq has enhanced sensitivity for detecting weaker yet specific CTCF-DNA interactions.

Analysis of motif composition revealed that 99.67% of the identified CTCF recognition sequences (118,008 of 118,395) contained the core CTCF motif-1 (Fig. 1, C and D). For downstream analyses, these sequences were classified into six categories based on combinations of motif-1 with flanking motifs. Approximately half of the CTCF-binding events (50.0%) involved at least one additional flanking motif (classes 1-5). We identified two upstream motifs, motif-2 and motif-2′, which differ primarily at the seventh and eighth nucleotide positions. In contrast, the downstream 8-mer motif-3 showed significantly higher consistency at the third and sixth positions. Only a small fraction of CTCF recognition sequences (0.33%) contained atypical motif configurations (fig. S1E).

Genomic distribution analysis revealed that promoter and exon regions were enriched for CTCF-binding sites relative to random expectation (two- to fourfold enrichment), whereas intronic and intergenic regions showed no such enrichment (Fig. 1E). Consistent with this observation, promoter and exon regions of transcriptionally active genes were more frequently occupied by CTCF than those of inactive genes (fig. S2, A and B), suggesting an association between CTCF binding and gene activity. Furthermore, CTCF sites located within 0-300 bp upstream of transcription start sites (TSSs) preferentially displayed forward motif orientation (Fig. 1F). This orientation bias was not explained by promoter activity per se but instead correlated with intrinsic CTCF-binding affinity (fig. S2, C to H).

### DeepCTCF enables prediction of quantitative CTCF-DNA binding affinity

Deep learning approaches have been widely applied to predict transcription factor binding (*43, 45*); however, most existing frameworks either generate binary binding predictions or are optimized for long genomic sequences. To quantitatively predict CTCF-binding affinity for short 42-bp DNA sequences, we developed a convolutional neural network (CNN), termed DeepCTCF, and systematically optimized the number of convolutional layers and filter sizes (Fig. 2A, and fig. S3, A and B).

**Fig. 2.**
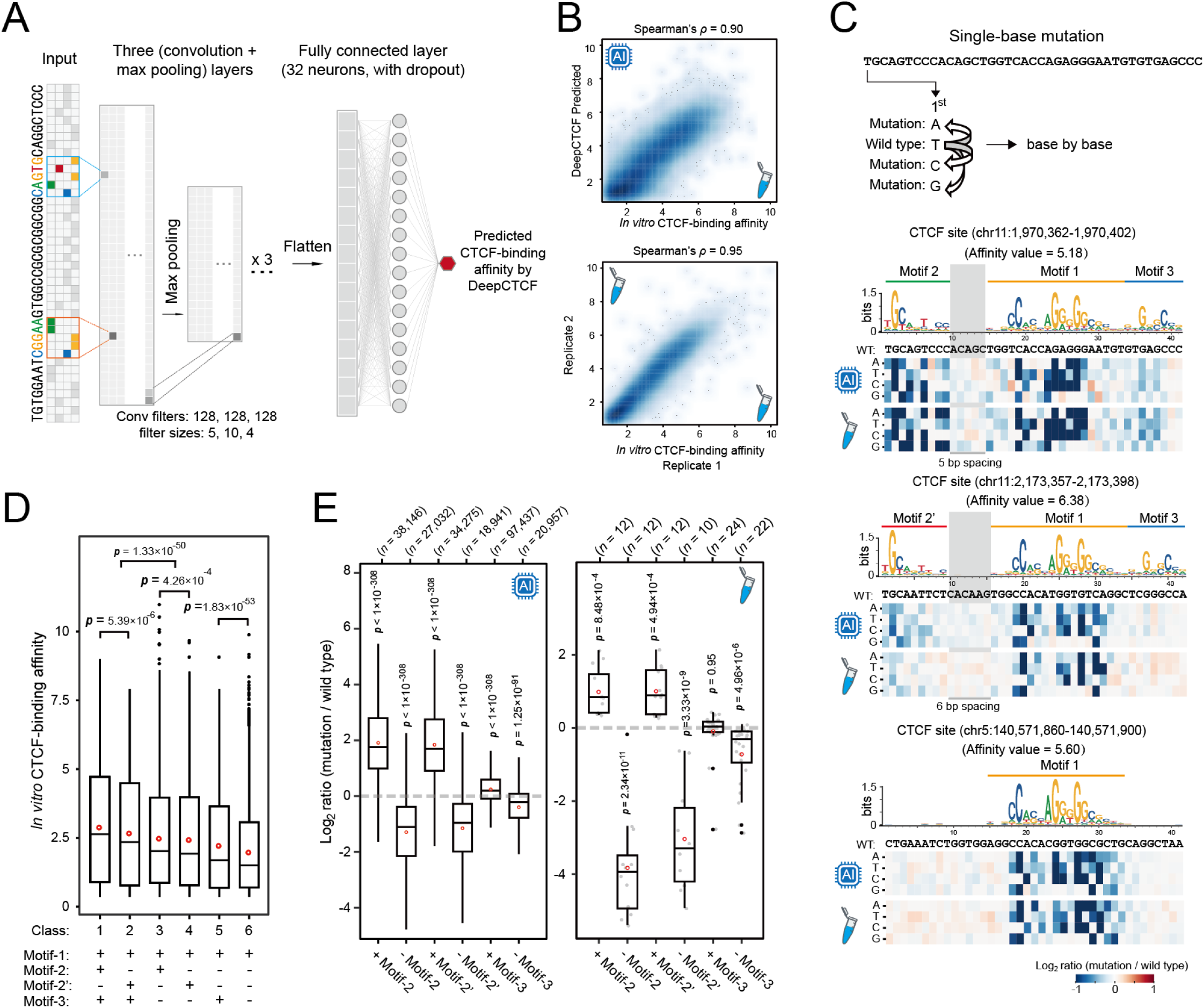
DeepCTCF quantitatively predicts CTCF-DNA binding affinity from sequence. (**A**) Architecture of the convolutional neural network (CNN) DeepCTCF, trained to predict CTCF-binding affinity directly from 42-bp DNA sequences. (**B**) Scatter plots comparing DeepCTCF-predicted and MpEMSA-seq-measured CTCF-binding affinity values (top), and the correlation between two biological replicates of MpEMSA-seq measurements (bottom). Color indicates local density. (**C**) Schematic of single-base-pair mutational analysis. Heat maps show log_2_ fold changes in CTCF-binding affinity upon single-nucleotide substitutions, as predicted by DeepCTCF (top) and measured experimentally (bottom). (**D**) Comparison of CTCF-binding affinity across the six classes of CTCF recognition sequences defined in Fig. 1C. (**E**) Effects of adding or removing sequence elements corresponding to motif-2, motif-2′, or motif-3 on CTCF-binding affinity. Box plots show log_2_ changes in affinity predicted by DeepCTCF (left) and measured experimentally (right). Statistical significance was assessed using an unpaired two-tailed Student’s *t* test.

DeepCTCF performance was evaluated using a held-out test set comprising one-third of the full dataset. Predicted affinity values showed a strong correlation with experimentally measured values (Spearman’s *ρ* = 0.90), approaching the concordance observed between biological replicates (*ρ* = 0.95) (Fig. 2B). We next benchmarked DeepCTCF against established prediction approaches. DeepCTCF substantially outperformed a position weight matrix (PWM)-based algorithm (*46*) (*ρ* = 0.33) as well as the CNN-based BPNet model (*43*) (*ρ* = 0.58), which was originally designed for predictions from 1-kb genomic sequences. Beyond quantitative prediction, DeepCTCF enables single-base-resolution inference of CTCF-binding affinity on a genome-wide scale. As an illustrative example, DeepCTCF accurately identified 27 CTCF-binding sites within the protocadherin alpha (Pcdhα) gene cluster and its associated regulatory regions (fig. S3C) (*6*).

A central question is how individual nucleotides within a CTCF recognition sequence contribute to overall binding affinity. To address this, we performed systematic single-base mutational analyses using both DeepCTCF predictions and experimental MpEMSA-seq measurements (Fig. 2C and table S2). Mutations within positions corresponding to CTCF motifs frequently led to reduced binding affinity (Fig. 2C, and fig. S4, A and B). Notably, high-affinity CTCF sites were generally less sensitive to single-base substitutions (fig. S4A), and an artificial high-affinity site exhibited minimal changes upon mutation (fig. S4C). In contrast, intermediate- and low-affinity sites showed substantially greater sensitivity to base substitutions (fig. S4B). Consistent with these observations, susceptibility to mutation exhibited a negative correlation with intrinsic CTCF-binding affinity (fig. S4D). Overall, DeepCTCF-predicted affinity changes were strongly correlated with experimentally measured values (Pearson’s *r* = 0.87) (fig. S4E).

We next compared binding affinity across different classes of CTCF recognition sequences (Fig. 2D). CTCF sites containing an upstream motif (motif-2 or motif-2′; classes 1-4) displayed significantly higher affinity than sites lacking an upstream motif (class 1/2 versus 5; class 3/4 versus 6). Similarly, CTCF-binding events incorporating the downstream motif-3 exhibited stronger affinity than those without this motif (1 versus 3; 2 versus 4; 5 versus 6).

To directly assess the contribution of flanking motifs to CTCF binding, we first used DeepCTCF to predict affinity changes resulting from the removal or introduction of sequences containing individual flanking motifs (Fig. 2E, left). We then synthesized oligonucleotides harboring newly introduced motif-2, motif-2′, or motif-3 sequences and measured their binding affinity using MpEMSA-seq (table S3). Relative to control sequences, the introduction of motif-2 or motif-2′ resulted in an average ∼2-fold increase in CTCF-binding affinity (Fig. 2E, right). Conversely, removal of any flanking motif significantly reduced binding affinity. Although mutations affecting motif-3 caused a more modest reduction (approximately 40% on average), disruption of motif-2 or motif-2′ led to a pronounced decrease in affinity, with an average reduction exceeding eightfold.

### Distinct sequence and spacing properties of upstream motifs 2 and 2′

The affinity values of CTCF sites differed significantly between classes 1 and 2, as well as between classes 3 and 4 (Fig. 2D), indicating that sites utilizing motif-2 generally exhibit higher binding affinity than those employing motif-2’. Consistent with this difference, the spacing between motif-2/2’ and motif-1 also varied between these classes (Fig. 3A, top panel).

**Fig. 3.**
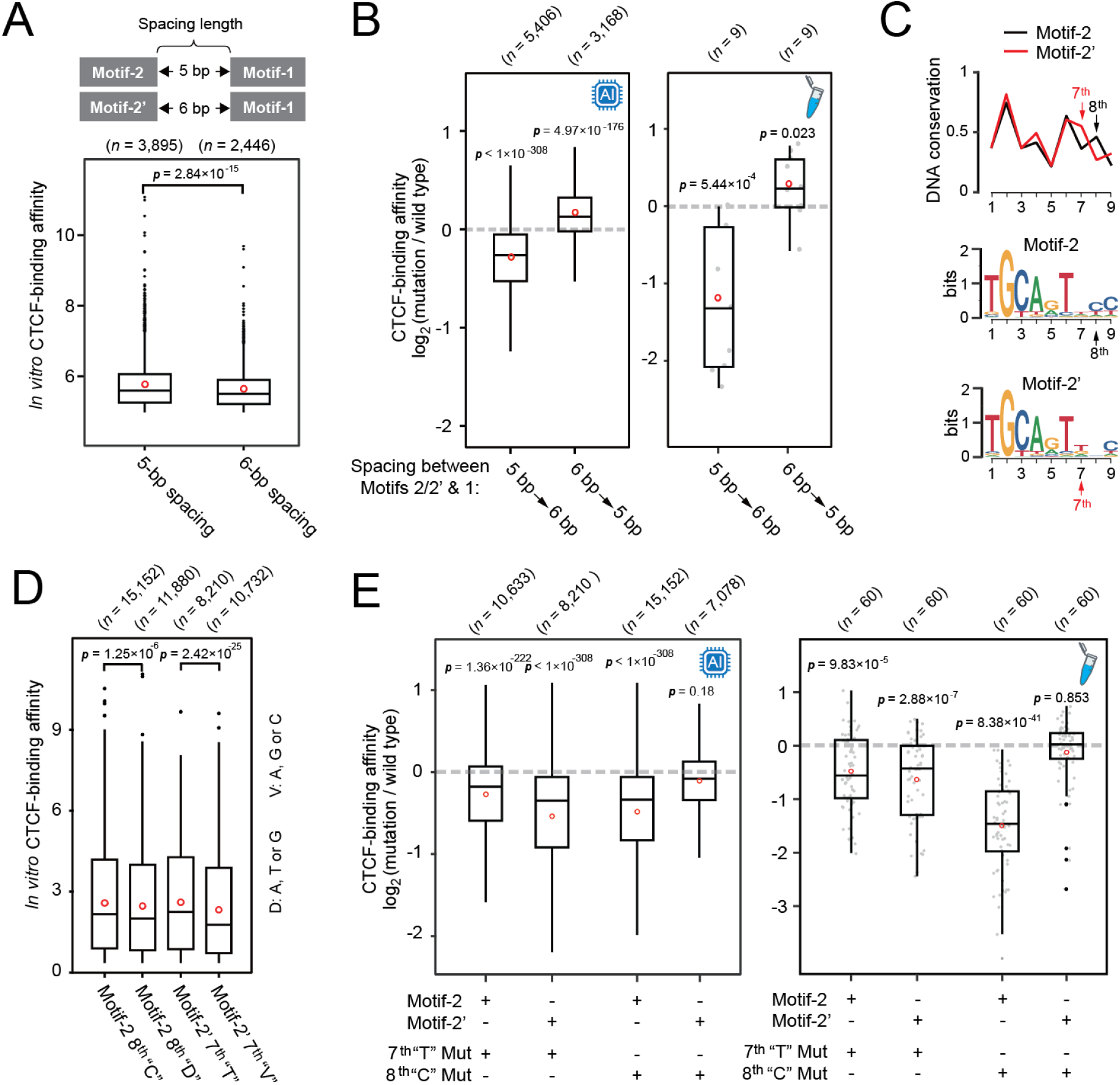
Spacing and base-specific constraints in motif-2 fine-tune CTCF binding affinity. (**A**) Schematic illustrating variable spacing between the upstream motif (motif-2 or motif-2′) and the core motif-1 (top). Box plots (bottom) compare CTCF-binding affinity for sequences containing motif-2 plus motif-1 versus motif-2′ plus motif-1. (**B**) Effects of spacing length between the upstream motif and motif-1 on CTCF-binding affinity, as predicted by DeepCTCF (left) and measured experimentally (right). To control for intrinsic sequence differences between motif-2 and motif-2′, all tested sequences contain “TC” at the seventh and eighth nucleotide positions. (**C**) Evolutionary conservation of motif-2 and motif-2′ across vertebrates. Conservation profiles are shown for motif-2 (middle) and motif-2′ (bottom), with the seventh and eighth nucleotide positions highlighted by red and black arrows, respectively. (**D**) Box plots compare CTCF-binding affinity for sequences carrying either T or C at the seventh or eighth nucleotide position. (**E**) Effects of single-nucleotide substitutions at the seventh or eighth positions on CTCF-binding affinity. Box plots show log_2_ changes in affinity relative to the corresponding wild-type sequences. Statistical analysis was performed using an unpaired two-tailed Student’s *t* test.

We therefore examined the effect of spacing length on CTCF binding affinity. Among the top 10,000 highest-affinity CTCF sites, 3,895 sites contained a 5-bp spacing between motif-2 and motif-1, whereas only 2,446 sites exhibited a 6-bp spacing. Moreover, sequences with a 5-bp spacing displayed significantly higher affinity than those with a 6-bp spacing (Fig. 3A, bottom panel).

To directly test the contribution of spacing length, we converted a set of 5-bp-spacing CTCF sites to 6-bp-spacing sites by inserting a single nucleotide into the spacer region, and reciprocally converted 6-bp-spacing sites to 5-bp by deleting one nucleotide (Fig. 3B). Transition from 5- to 6-bp spacing resulted in a marked reduction in CTCF binding affinity, whereas conversion from 6- to 5-bp spacing led to a significant increase (table S4), demonstrating a causal role for spacing length in regulating CTCF binding strength.

We next compared the evolutionary conservation of motif-2 and motif-2’. Vertebrate conservation analysis revealed that the 7^th^ base of motif-2’ is more conserved, whereas the 8^th^ base of motif-2 shows higher conservation on average (Fig. 3C and fig. S5A).

Correspondingly, CTCF sites containing a “C” at the 8^th^ position of motif-2 exhibited significantly higher affinity than those lacking this base, while sites containing a “T” at the 7^th^ position of motif-2’ also showed elevated affinity (Fig. 3D). These effects were independently supported by AI-based affinity prediction (Fig. 3E, left panel).

To experimentally validate these observations, we mutated the 7^th^ “T” in motif-2’ and the 8^th^ “C” in motif-2 in a set of representative sequences (table S5). Mutation of the 7^th^ “T” resulted in a moderate reduction in binding affinity (approximately 30% on average). In contrast, mutation of the 8^th^ “C” caused a pronounced decrease in affinity for sequences containing motif-2 (an average reduction of 67%), but had little effect on sequences carrying motif-2’ (Fig. 3E, right panel). Consistently, CTCF sites harboring a “T” at the 7^th^ position of motif-2 exhibited significantly higher affinity than those lacking this base (fig. S5B), highlighting a critical role of this nucleotide in strengthening CTCF binding.

### High G/C content in motif-1 flanking regions suppresses CTCF binding

Although CTCF recognition sequences can be grouped into six motif combinations (Fig. 1C), substantial variability in binding affinity persists within each class (Fig. 2D), implying that additional sequence features beyond motif composition contribute to quantitative CTCF-binding strength. To identify regulatory features beyond the core motif-1, we systematically interrogated the contribution of flanking sequences to CTCF binding. All possible upstream 13-/14-bp or downstream 10-bp sequences were generated and individually substituted for the corresponding regions of a reference CTCF site (Fig. 4A, top panel). The binding affinity of each resulting sequence was then predicted using DeepCTCF (Fig. 4A, bottom panel).

**Fig. 4.**
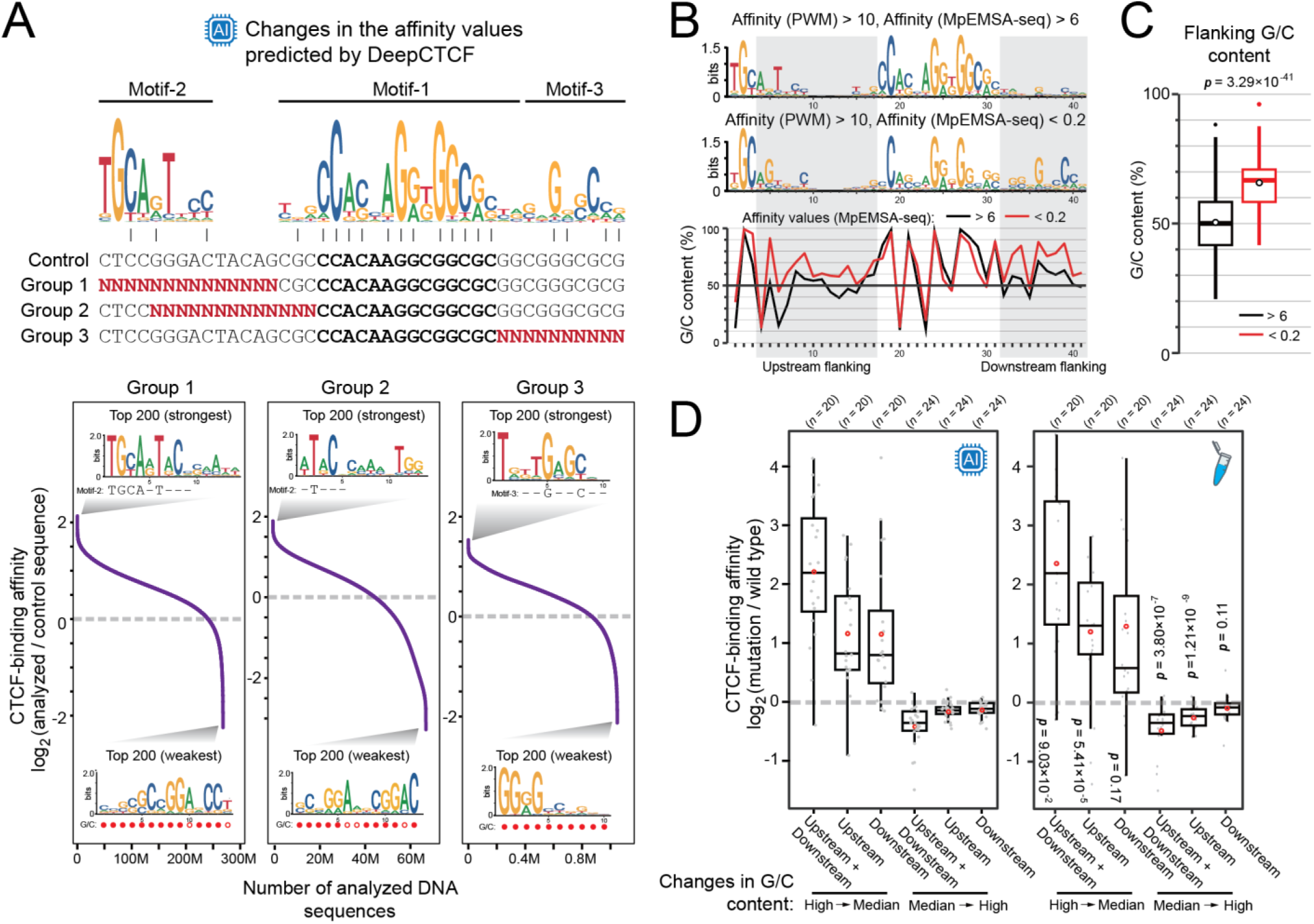
Elevated G/C content in key motif-1 flanking sequences inhibits CTCF binding. (**A**) DeepCTCF-predicted CTCF-binding affinity for all possible upstream (groups 1 and 2) or downstream (group 3) sequences combined with a fixed sequence matching the CTCF core motif-1. In the bottom panel, affinity values for each sequence are normalized to a control sequence. Sequence logos show consensus motifs derived from the top 200 sequences with the strongest or weakest predicted affinity. (**B**) Sequence features associated with large differences in CTCF-binding affinity. The top panel shows consensus motifs of sequences identified by a position weight matrix (PWM)-based scan (score > 10). The bottom panel shows G/C content profiles for sequences with high affinity (> 6; black) or low affinity (< 0.2; red). Shaded regions indicate flanking positions adjacent to key motif-1. (**C**) Effects of altering G/C content within key motif-1 flanking sequences on CTCF-binding affinity. Box plots show changes in affinity relative to the corresponding control sequences. Statistical significance was assessed using a two-sided unpaired Student’s *t* test.

Among the top 200 sequences with the highest predicted affinity, the upstream substitutions in groups 1 and 2 were strongly enriched for motifs resembling motif-2, whereas a motif related to motif-3 was identified in the top-ranking downstream substitutions (group 3). Unexpectedly, consensus sequences also emerged among the bottom 200 sequences with the weakest predicted affinity. These low-affinity consensus sequences were characterized by an excessively high guanine and cytosine (G/C) content, and their predicted binding strength was reduced by more than 75%.

Consistently, CTCF sites carrying identical core motifs (PWM score > 10) nonetheless exhibited a wide range of binding affinities (Fig. 4, B and C, and fig. S6). Notably, sequences with the weakest CTCF binding (affinity < 0.2) showed significantly higher G/C content in the flanking regions of key motif-1 compared to sequences with strong binding (affinity > 6), indicating that flanking base composition can negatively modulate CTCF binding.

To directly test this effect, we experimentally altered the G/C content of motif-1 flanking regions at non-consensus positions, converting high G/C content to an intermediate level (∼50%), or vice versa (Fig. 4D and table S6). Modifying both upstream and downstream flanking regions produced the strongest effects. Reducing G/C content from high to intermediate levels increased CTCF-binding affinity by more than twofold on average, whereas increasing G/C content from intermediate to high decreased binding strength. Together, these results demonstrate that elevated G/C content in motif-1 flanking sequences acts as a negative determinant of CTCF binding affinity.

### Disease-associated variants modulate CTCF-binding affinity

Most disease-associated variants identified by genome-wide association studies (GWAS) reside in noncoding regions of the genome, including CTCF-binding sites (*47, 48*). We therefore systematically evaluated the impact of known human genetic variants on CTCF binding using DeepCTCF. This analysis identified more than 1.2 million variants predicted to significantly alter CTCF-binding affinity (log_2_ fold change > 1 or < -1) (Fig. 5A, and fig. S7, A and B).

**Fig. 5.**
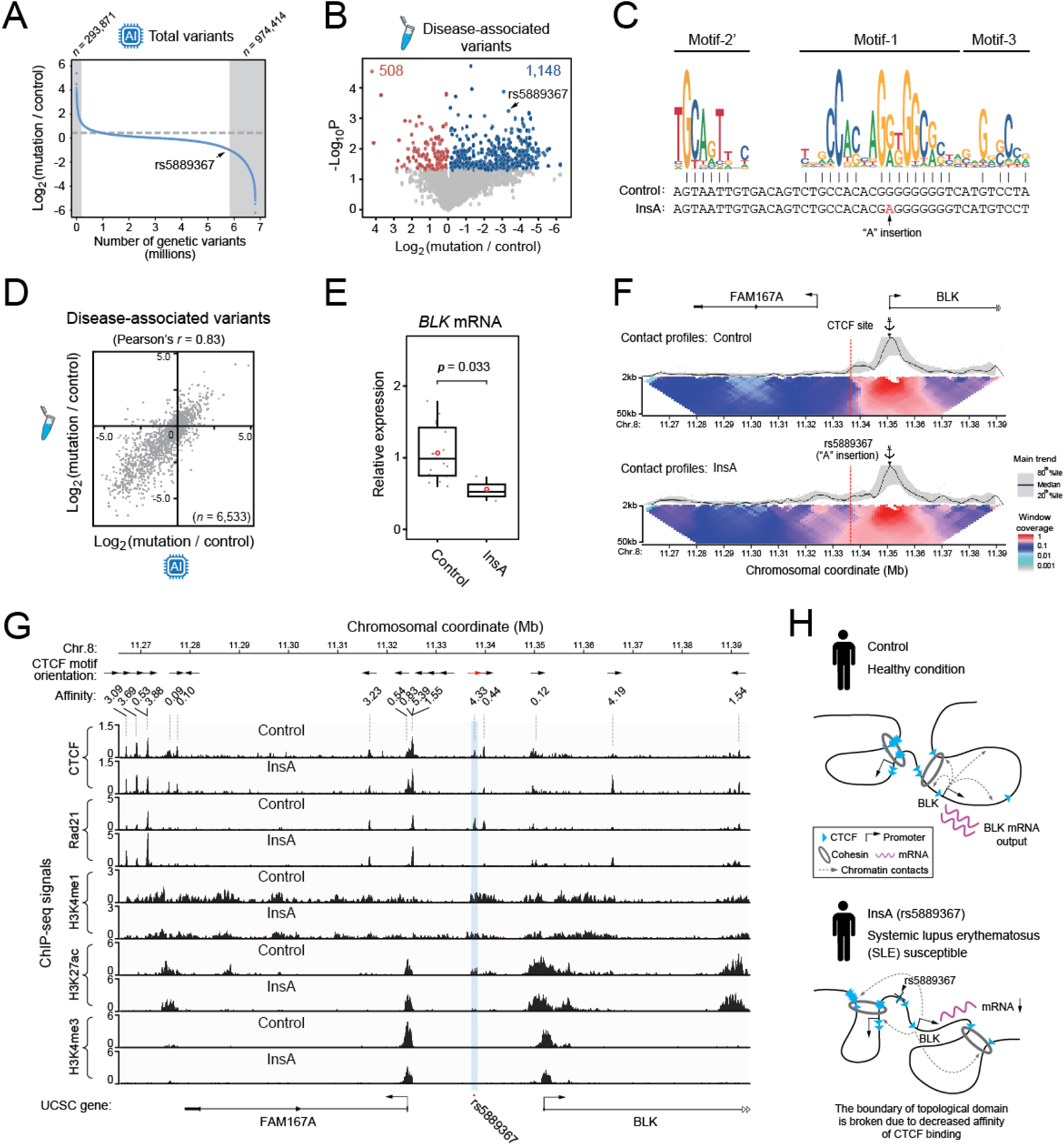
Identification and functional validation of disease-associated variants that alter CTCF-binding affinity. (**A**) DeepCTCF-predicted changes in CTCF-binding affinity for DNA sequences carrying genetic variants. Variants from the latest dbSNP release were ranked by log_2_ (mutant/control) affinity ratios; variants with log_2_ fold changes greater than 1 or less than -1 are highlighted (gray shading). (**B**) Volcano plot showing CTCF sites containing disease-associated variants with statistically significant changes in binding affinity. Sites with significantly decreased affinity (*p* < 0.05) are shown in blue, and those with increased affinity in red. The CTCF site associated with SNP rs5889367 is indicated by a black arrow. *P* values were calculated using a paired two-tailed Student’s *t* test. (**C**) Sequences of the control allele and the InsA variant shown relative to the CTCF motifs. The inserted adenine is highlighted in red. (**D**) Correlation between DeepCTCF-predicted and experimentally measured changes in CTCF-binding affinity for 6,533 disease-associated variants. Each gray dot represents an individual variant. (**E**), Expression levels of the *BLK* gene in control and InsA cells. Statistical analysis was performed using the unpaired two-tailed Student’s *t* test. (**F**) Chromatin contact profiles of the *BLK* promoter in control (top) and InsA (bottom) cells. Black lines indicate median interaction frequencies computed in 5-kb windows, with gray bands denoting the 20^th^-80^th^ percentiles. Heat maps below show median normalized contact intensities across window sizes ranging from 2 to 50 kb. The dashed red line marks the CTCF site associated with rs5889367, and the black anchor indicates the viewpoint position. Chr., chromosome. (**G**) Genomic binding profiles of CTCF, the cohesin subunit RAD21, and histone marks H3K4me1, H3K27ac, and H3K4me3. Signal intensities are shown as reads per million. CTCF motif orientations are indicated by arrows. The region encompassing rs5889367 is highlighted in blue, and the associated CTCF site is indicated by a red arrow. (**H**) A model illustrating how rs5889367 reduces *BLK* expression by weakening CTCF-binding affinity and disrupting a topological domain boundary, thereby contributing to susceptibility to systemic lupus erythematosus (SLE).

To experimentally evaluate the predictive performance of DeepCTCF, we assayed a subset of 6,533 disease-associated genetic variants using MpEMSA-seq (fig. S7, C and D, and table S7). Among these variants, 508 resulted in enhanced CTCF-binding activity, whereas 1,148 caused a significant reduction in CTCF-binding affinity. Notably, rs5889367, a single-nucleotide polymorphism associated with systemic lupus erythematosus (SLE) and located near the SLE risk gene *BLK* (*49-51*), was validated to markedly reduce CTCF binding by introducing an adenine into the core motif-1 (Fig. 5C). Changes in CTCF-binding affinity predicted by DeepCTCF were strongly correlated with experimentally measured binding strengths (Pearson’s *r* = 0.83; Fig. 5D), supporting the accuracy and robustness of DeepCTCF for quantitative functional screening of noncoding variants.

To validate the functional impact of rs5889367 in a cellular context, we precisely inserted an adenine at the endogenous CTCF site in human B lymphocytes (Raji cells) using prime editing. Three independent clonal cell lines carrying the insertion (InsA, clones 1-3) and three matched control clones lacking the insertion (Control, clones 1-3) were isolated. The adenine insertion caused a pronounced loss of CTCF binding at the edited site and led to an approximately 50% reduction in *BLK* mRNA levels (Fig. 5E and fig. S7E). Chromatin conformation capture (4C-seq) revealed increased interactions between the *BLK* promoter and multiple upstream loci in InsA cells, indicative of impaired insulation (Fig. 5F).

Similar interaction changes were observed using an independent viewpoint within the *BLK* promoter (fig. S7F). Consistently, ChIP-seq analysis demonstrated a near-complete loss of CTCF and cohesin subunit Rad21 binding at the rs5889367-associated site in InsA cells, accompanied by alterations in local chromatin states marked by H3K4me1, H3K27ac, and H3K4me3 (Fig. 5G). Given that this CTCF site is oriented in the forward direction and is predicted to act as a boundary for the downstream topological domain, the adenine insertion effectively disrupted domain insulation. Accordingly, chromatin interactions between this CTCF site and its downstream intradomain regions were significantly reduced (fig. S7, G and H).

Together, these results support a model in which rs5889367 compromises CTCF binding, disrupts topological domain organization, and consequently down-regulates *BLK* expression, thereby contributing to SLE susceptibility (Fig. 5H).

## Discussion

Herein, we quantitatively determine the strength of CTCF binding to hundreds of thousands of unmodified DNA sequences derived from the human genome by a high-throughput biochemical assay, MpEMSA-seq. Compared to the methods using genomic DNA (*44, 52*), the sequence complexity in our assay is reduced by over two hundred fold, enabling more precise assessment of CTCF-binding strength across diverse sequences. We demonstrate that CTCF can specifically bound to at least 117,974 genomic sequences covering more than 1‰ of the genome. The number of identified CTCF sites is approximately 4-fold higher than that obtained using the previous method. Thus, our method provides, to our knowledge, the most comprehensive quantitative map of *in vitro* CTCF-binding affinity across the human genome to date. Based on the large, high-quality dataset, we further develop a deep-learning model, DeepCTCF, to directly predict the strength of CTCF binding from DNA sequence.

We use a complementary combination of DeepCTCF and MpEMSA-seq to investigate how DNA sequence features influence binding affinity to CTCF. We show that, almost all (99.6%) CTCF-binding sites contain the core motif-1, while CTCF sites employing one or two flanking non-core motifs exhibit on average stronger CTCF-binding ability. We speculate that the two highly similar upstream motifs, 2 and 2’, may share a common evolutionary origin, yet contribute differently to CTCF-binding strength through distinct sequence-dependent mechanisms. First, the spacing between motif-2/2’ and motif-1 emerges as a critical determinant, with a 5-bp spacing arrangement conferring substantially stronger binding than a 6-bp spacing. Second, the 8^th^ base “C” of motif-2 can further increase CTCF-binding affinity relative to that of motif-2’. Beyond consensus motifs, we reveal that nucleotide composition at non-consensus positions within CTCF-binding sites can also exert a pronounced effect on binding strength.

Together, our findings identify multiple layers of sequence information that bidirectionally regulate CTCF-binding affinity. Specifically, we define three key principles: (i) the quantitative contribution of flanking motifs to CTCF binding, (ii) the mechanistic distinction between two upstream motif variants, and (iii) the inhibitory effect of high G/C content in critical motif-1 flanking sequences. Based on these principles, DeepCTCF enables rational exploration and engineering of CTCF recognition sequences with tunable binding strength, extending beyond binary binding predictions.

In this study, we further apply DeepCTCF to predict binding affinity changes for more than 6 million human genetic variants. We experimentally verify 6,533 disease-associated variants, revealing a strong correlation between predicted and experimentally measured CTCF-binding strength. These results support the quantitative accuracy and generalizability of DeepCTCF across diverse sequence contexts. Collectively, our framework establishes a direct and predictive link between DNA sequence variation and CTCF-binding affinity, providing a powerful tool for investigating how regulatory sequence alterations may contribute to disease mechanisms and chromatin architectural defects.

While this study establishes a quantitative framework for understanding how DNA sequence encodes intrinsic CTCF-binding affinity, several limitations define the scope of the present results and inform their appropriate interpretation. The affinity measurements and models reported here were obtained under controlled *in vitro* conditions and are therefore designed to capture the sequence-encoded component of CTCF-DNA recognition in isolation. As a consequence, the reported affinity should be viewed as a biochemical baseline for CTCF binding rather than a direct measure of regulatory activity or occupancy in living cells. *In vivo*, CTCF binding and function are further shaped by multiple layers of regulation that are not incorporated into the current framework.

Notably, DNA methylation has been shown to modulate CTCF occupancy at specific genomic loci, and systematic *in vitro* measurements incorporating defined cytosine methylation states will be required to quantify its effects on binding affinity. In addition, chromatin accessibility, nucleosome positioning, transcription, and interactions with cofactors are expected to modulate how intrinsic sequence affinity is translated into functional CTCF occupancy. Future efforts that integrate quantitative sequence-derived affinity measurements with chromatin context and cellular regulatory information will be essential for applying the present framework to predict CTCF occupancy and function *in vivo*.

## Materials and Methods

### Experimental Design

### Cell Culture

HEK293T cells were cultured in Dulbecco’s modified Eagle’s medium (DMEM; Gibco, 11995) supplemented with 10% (v/v) fetal bovine serum (FBS; Royacel, RYS-KF22) and 1% penicillin–streptomycin (Gibco, 15140). Raji cells were maintained in RPMI 1640 medium (Gibco, 11875) supplemented with 10% (v/v) FBS and 1% penicillin-streptomycin. All cells were cultured at 37 °C in a humidified incubator with 5% CO_2_.

### MpEMSA-seq

DNA probes for multiple parallel electrophoretic mobility shift assay followed by high-throughput sequencing (MpEMSA-seq) were generated by polymerase chain reaction (PCR) using customized oligonucleotide pool (GenScript) as template. To precisely measure intrinsic CTCF-binding strength, the length of all tested CTCF-sites was uniformly fixed at 42 bp. The DNA-binding domain of CTCF were synthesized *in vitro* using the cell-free TNT T7 Quick Coupled Transcription/Translation System (Promega; L1170).

EMSA experiment was then performed using the LightShift Chemiluminescent EMSA kit (Thermo; 20148) according to previously described procedures (*9*). Briefly, DNA probes were incubated with the *in vitro-*translated proteins in the binding buffer containing 10 mM Tris, 50 mM KCl, 1 mM dithiothreitol, 2.5% (v/v) glycerol, 5 mM MgCl_2_, 50 ng/μl poly (dI-dC), 0.1% (v/v) Nonidet P-40 (NP-40), and 0.1 mM ZnSO_4_ at room temperature for 20 minutes.

Samples were electrophoresed in ice-cold 0.5×TBE buffer (Sangon; B040124) at 100V for 85 minutes on 5% nondenaturing polyacrylamide gels. Gel bands corresponding to the DNA-protein complexes, detected by chemiluminescence, were excised. DNA was purified by phenol-chloroform extraction followed by ethanol precipitation.

Amplification of the MpEMSA-seq library for Illumina sequencing was performed by PCR using primers containing Illumina adapter sequences. Raw reads were obtained by high-throughput sequencing, and uniquely mapped reads were counted for each CTCF site and normalized to one million total reads for the shifted (RPM^shift^) and input (RPM^input^) libraries, respectively. The relative binding affinity of CTCF for each tested site was calculated as the ratio RPM^shift^ / RPM^input^.

### DeepCTCF

The DeepCTCF model is based on a convolutional neural network (CNN) architecture that takes one-hot-encoded 42-bp DNA sequences (A = [1,0,0,0], T = [0,0,0,1], G = [0,0,1,0], C = [0,1,0,0]) as input features to predict the quantitative affinity of CTCF binding. DeepCTCF consists of three one-dimensional convolutional layers (number of filters = 128, 128, and 128; kernel sizes = 5, 10, and 4, respectively), each followed by a rectified linear unit (ReLU) activation and max-pooling (pool size = 2).

Following the convolutional layers, a fully connected layer with 32 neurons was applied, followed by a ReLU activation and dropout (rate = 0.2). The DeepCTCF architecture was optimized by systematically varying the number of convolutional layers and convolutional filter sizes (fig. S3, A and B).

Model performance was evaluated using the Spearman rank correlation coefficient (*ρ*) on an independent held-out test set. DeepCTCF was implemented and trained using Keras (v2.4.3) with TensorFlow (v2.4.1) as the backend, using mean squared error (MSE) as the loss function and early stopping with a patience of 10 epochs. Model parameters were optimized using the Adam optimizer (learning rate = 1 ×10^-4^) for up to 50 epochs, with a batch size of 32 sequences.

To investigate the effects of human genetic variants on CTCF-binding affinity, the latest release of genetic variants was downloaded from the dbSNP database (GCF_000001405.40). For each variant, the affinity values of CTCF recognition sequences carrying the variant allele (mutation) and the corresponding reference sequence (control) were predicted using DeepCTCF, and the relative effect of the variant was quantified as log_2_ (mutation/control).

### RNA isolation and qRT-PCR

Cells for qRT-PCR analysis were harvested in TRIzol Reagent (Thermo; 15596018CN). The quantity of RNA was determined by a NanoDrop-2000 spectrophotometer (NanoDrop). Total RNA was reverse-transcribed into cDNAs using the HiScript II Q RT SuperMix reagents (Vazyme; R223-01). Real-time quantitative polymerase chain reaction was performed using the SYBR qPCR Master Mix reagents (Vazyme; Q712-02) on a QuantStudio Real-Time PCR system (Applied Biosystems). Fold differences were calculated using the △△*C*_t_ method.

### Prime editing

The prime editing strategy was used to insert a single adenine nucleotide into the CTCF recognition sequence in Raji cells. The prime editing guide RNA (pegRNA) was designed using an online tool (http://pegfinder.sidichenlab.org/) (*53*).

To assemble the nicking guide RNA (ngRNA) expression vector, the pKLV-U6gRNA (BbsI)-PGKpuro2ABFP plasmid (Addgene; #50946) was digested using BbsI (NEB; R3059L). A pair of oligonucleotides corresponding to the ngRNA protospacer was annealed, and ligated into the BbsI-linearized plasmid. The inserted sequence was verified by Sanger sequencing.

For construction of the pegRNA expression vector, the pKLV-U6gRNA (BbsI)-PGKpuro2ABFP plasmid was digested with both BbsI and BamHI (NEB; R0136S). The pegRNA components, including guide RNA oligonucleotides, gRNA scaffold oligonucleotides, reverse transcription (RT) template, and prime binding site oligonucleotides, were assembled into the double-digested plasmid. The assembled pegRNA sequence was confirmed by Sanger sequencing.

For prime editing, 2×10^6^ cells were harvested and washed with PBS prior to electroporation. For transfection, 10 μg of the pCMV-PE2-P2A-GFP plasmid (Addgene; #132776), 10 μg of pegRNA plasmid, and 5 μg of ngRNA plasmid were mixed and delivered into cells using a Neon transfection system (Thermo Fisher Scientific) under the following conditions: 1,400 V, 10 ms, and three pulses.

After transfection, cells were plated into a 6-well plate and incubated for 72 hours. Single cells displaying strong GFP and BFP fluorescence were sorted by fluorescence-activated cell sorting (FACS) into a 96-well plate. After 14 days of clonal expansion, the genotypes of individual subclones were validated by Sanger sequencing.

### ChIP-qPCR and ChIP-seq

Ten million cells were cross-linked at room temperature for 10 minutes with 1% (v/v) formaldehyde (Thermo; 28908), followed by quenching with glycine (Sigma; G5417) to a final concentration of 0.125 M. The cross-linked cell pellet was resuspended in 1 mL of ice-cold RIPA Buffer (10mM Tris pH7.5; 0.15M NaCl; 1% Triton X-100; 1mM EDTA; 0.1% Sodium deoxycholate; 0.1% SDS; 1×Roche EDTA-free protease inhibitor cocktails), and incubated with gentle rotation at 4°C for 15 minutes. This lysis step was repeated once, after which the isolated nuclei were resuspended in sonication buffer (10 mM Tris–HCl, pH 7.5; 150 mM NaCl; 1% Triton X-100; 1 mM EDTA; 0.1% sodium deoxycholate; 1% SDS; 1×Roche EDTA-free protease inhibitor cocktail).

Chromatin was sheared using a Scientz 08-III sonicator (Scientz) with 90% amplitude for a total of 25 minutes (25 cycles of 30 seconds on and 30 seconds off). The supernatant was collected and diluted 10-fold in dilution buffer (10 mM Tris-HCl, pH 7.5; 150 mM NaCl; 1% Triton X-100; 1 mM EDTA; 0.1% sodium deoxycholate; 1×Roche EDTA-free protease inhibitor cocktail).

For ChIP, chromatin lysates were incubated with anti-CTCF antibody (Abcam; ab70303) overnight at 4 °C with gentle rotation. DNA-protein complexes were captured using protein G magnetic beads (Thermo; 10004D). The beads were sequentially washed once each with RIPA buffer containing 0.15 M NaCl, RIPA buffer containing 0.4 M NaCl, RIPA buffer without NaCl, LiCl wash buffer (0.25 M LiCl; 0.5% NP-40; 0.5% sodium deoxycholate; 1×Roche EDTA-free protease inhibitor cocktail), and 1×Tris-EDTA buffer.

After reverse crosslinking, DNA was purified by phenol-chloroform-isoamyl alcohol extraction. DNA concentration was determined using a Qubit dsDNA HS Assay Kit (Thermo; Q32854) on a Qubit fluorometer (Thermo).

For ChIP-qPCR, purified DNA was quantified by real-time quantitative PCR using FastStart Universal SYBR Green Master Mix (Roche; 4913914001) on a qTOWER3G Touch Real-Time PCR system (Analytik Jena). All qPCR reactions were performed in triplicate. Relative enrichment was calculated based on a standard curve generated from serially diluted input DNA (chromatin without antibody), and values were normalized to a control genomic region. All ChIP-qPCR data were obtained from two or three independent biological replicates. Primer sequences used for ChIP-qPCR are listed in table S8.

For ChIP-seq, libraries were prepared following the same ChIP procedure using the NEBNext DNA Library Prep Kit (NEB; E7645) according to the manufacturer’s instructions. Antibodies used for ChIP-seq included anti-CTCF (Abcam; ab70303), anti-Rad21 (Abcam; ab992), anti-H3K4me1 (Abcam; ab8895), anti-H3K27ac (Abcam; ab4729), and anti-H3K4me3 (Abcam; ab8580).

After Illumina sequencing, reads were aligned to the GRCh37/hg19 reference genome using Bowtie2 (v2.3.5.1). SAMtools (v1.9) was used to filter and sort aligned reads, and PCR duplicates were removed using Picard (https://github.com/broadinstitute/picard). Peak calling was performed using MACS2 (v2.1.0) with the parameter “--SPMR -q 0.05”.

### 4C-seq

Five million cells were cross-linked at room temperature for 10 minutes with 1% (v/v) formaldehyde, and crosslinking was quenched by adding glycine to a final concentration of 0.125 M. Cell pellet was resuspended in ice-cold NP-40 lysis buffer (10mM Tris pH7.5, 0.15M NaCl, 0.5% NP-40, 1×Roche EDTA-free protease inhibitor cocktails) and incubated at 4°C with gentle rotation for 15 minutes. This lysis step was repeated once to isolate nuclei.

Nuclei were incubated in SDS permeabilization buffer (1X NEB Buffer 2 supplemented with 0.5% SDS) at 62°C for 7.5 minutes, after which chromatin was digested with MboI (NEB; R0147) at 37 °C for ∼12 h. The reaction was terminated by incubation at 65 °C for 20 minutes, and proximity ligation was performed using T4 DNA ligase (NEB; M0202).

Following reverse crosslinking, DNA was purified by phenol-chloroform-isoamyl alcohol extraction, followed by sodium acetate/ethanol precipitation. DNA concentration was measured using a Qubit dsDNA HS Assay Kit (Thermo Fisher Scientific). Subsequently, 25 μg of purified DNA was digested overnight at 37 °C with NlaIII (NEB; R0125). NlaIII was inactivated at 65 °C for 25 minutes, and the digested DNA fragments were diluted and self-circularized using T4 DNA ligase.

4C libraries were generated by inverse PCR using KOD-Plus-Neo DNA polymerase (TOYOBO; KOD-401) with primers listed in table S8. 4C-seq data was analyzed using the 4Cseqpipe (version 0.7) with the parameter: “-stat_type median, -trend_resolution 2000” (*54*).

### Statistical Analysis

All statistical analyses were performed using standard statistical methods, as indicated in the figure legends or main text. The number of participants (*n*), Pearson’s *r*, Spearman’s *ρ*, and exact *p* values are reported in the corresponding figures, legends or results sections.

## Acknowledgments

We thank Roger D. Kornberg (Stanford University), James W.D. King (Imperial College London), and Andrew Dimond (Imperial College London) for helpful discussions and comments.

## Funding

National Natural Science Foundation of China (32270570)

Shanghai Pudong New Area Health Commission grant (2024-PWXZ-06)

National Natural Science Foundation of China (32141004).

Science and Technology Commission of Shanghai Municipality (21DZ2210200).

H.F. and Y.X. were supported by the Zhiyuan Future Scholar Program.

## Author contributions

Conceptualization: Z.Y., Y.W., H.F., Q.W., N.S., and Y.G.;

Methodology: Z.Y., Y.W., G.H., Y.C., H.F., and Y.G.;

Formal analysis: Z.Y., Y.W., G.H., H.F., Q.W., N.S., and Y.G.;

Investigation: Z.Y., Y.W., G.H., Y.C., H.F., J.Z., Y.X., R.W., Z.Z., T.W., H.H., S.Q.X., W.W., W.G., Q.W., N.S., and Y.G.;

Resources: Z.Y., and Y.G.;

Writing – original draft: Z.Y., Y.W., G.H., N.S., and Y.G.;

Writing – review and editing: Z.Y., Y.W., G.H., Y.C., H.F., J.Z., Y.X., R.W., Z.Z., T.W., H.H., S.Q.X., W.W., W.G., Q.W., N.S., and Y.G.;

Funding acquisition: Y.G., N.S., Q.W., and W.G.;

Supervision: Y.G., N.S., Q.W., and W.G.

## Competing interests

Authors declare that they have no competing interests.

## Data and materials availability

The deep-learning model DeepCTCF, including trained weights and scripts required to reproduce the analyses, is available at https://github.com/Yin-Zihang/DeepCTCF. All materials used in this study are available upon reasonable request and are not subject to material transfer agreements.

## Notes

### Competing Interest Statement

The authors have declared no competing interest.

